# Combining environmental DNA and traditional sampling to assess the role of kelp aquaculture as an artificial habitat in Norway

**DOI:** 10.1101/2025.10.02.679966

**Authors:** Adam Jon Andrews, Hartvig Christie, Kim Præbel, Melissa Brandner, Ragnhild Grimm Torstensen, Larisa Lewis, Ragnhild Beck Hestness, Paul Ragnar Berg, Trine Bekkby, Kasper Hancke

## Abstract

Kelp aquaculture is expanding rapidly in the temperate Atlantic, yet its ecological role as an artificial habitat remains poorly understood. This study assessed the biodiversity associated with wave-exposed kelp farms in coastal Norway using both environmental DNA (eDNA) and traditional sampling methods. Fish and invertebrate communities were surveyed at two kelp farms and compared to nearby natural kelp forests and pelagic control sites. Results from gill-net fishing, camera observations, and invertebrate sampling revealed lower species richness and abundance at kelp farms compared to natural habitats. eDNA analyses (12S and CO1 markers) showed few differences in community composition between sites, and no species were identified as indicators of kelp farms. Traditional sampling detected the presence of the amphipod *Caprella mutica* in the kelp farms, indicating that operators can play roles in alien species spread. Overall, kelp farms supported distinct but less diverse communities than natural kelp forests, being more similar to pelagic control sites. Finally, eDNA provided limited insights compared to traditional methods. These findings suggest that kelp farms currently provide limited habitat provisioning in Norway, likely as a result of how and where kelp aquaculture is currently performed. We suggest the need for population-level assessments for relevant species like Atlantic lumpfish (Cyclopterus lumpus) and locally based assessments in collaboration with authorities to define nature-positive or negative outcomes to balance ecological and economic goals in the growing seaweed aquaculture industries of the Atlantic.

## 1. INTRODUCTION

In line with the UN Sustainable Development Goals, there is increasing interest in cultivated kelp as a low-trophic-level food source that is rapidly expanding beyond Asia (Duarte et al., 2021). However, there is little understanding of the environmental effects of kelp farming in new operating areas like the temperate Atlantic. Potential positive and negative effects are numerous, including plastic pollution, carbon sequestration, bioremediation, hosts for disease etc. (Campbell et al., 2019; Forbes et al., 2022; Theuerkauf et al., 2022). One key expected effect is habitat provision and ecological function that is a priority to assess given the global, European and national goals to protect and restore degraded marine habitats and populations like the Kunming–Montreal Global Biodiversity Framework targets as well as commercial interests in contributing to nature positivity (Duarte et al., 2021; Forbes et al., 2022; Pardo et al., 2025; WWF, 2025). These effects and goals create a strong need to identify the role of kelp aquaculture as a novel artificial habitat in the Atlantic.

Recent work has theorised the benefits of habitat provision of kelp farms as a potential aid towards the current biodiversity crisis (Theuerkauf et al., 2022), but there are few empirical studies conducted in the Atlantic to date. These include studies who found increases in the diversity and abundance of fishes compared to control sites in the south of England (Corrigan et al., 2024b) and conversely no evidence for this off Maine (Schutt et al., 2023). Similarly a study on benthic fish assemblages found no effects of kelp farms in Sweden (Visch et al., 2020a). Prey availability for fishes is apparent however; two preliminary studies found diverse epifauna communities at kelp farms, albeit in lower diversity and abundance than in natural kelp forests, with key differences such as greater abundances of amphipods at farm locations (Bekkby et al., 2023; Corrigan et al., 2024a) in agreement with an Irish study who noted modified epifauna communities in cultivated kelp (Walls et al., 2016). The potential for a modified epifauna community containing alien marine species is also apparent and may create ecological risks (Bernard et al., 2019). Increases in biodiversity at temperate kelp farms are expected given this is the case in the tropics (Theuerkauf et al., 2022) and that natural temperate kelp habitats are highly productivity environments, playing important roles for fishes including as nursery habitats (Steneck et al., 2002; Norderhaug et al., 2005; Christie et al., 2022).

Increased biodiversity is not necessarily positive, especially for non-native or not locally present species, given that they may alter food webs (Forbes et al., 2022) and modify the ecological function of important, often threatened and degraded, habitat types where they operate, such as kelp forests, seagrass meadows or soft sediments (Campbell et al., 2019; Theuerkauf et al., 2022). Meanwhile there are doubts on the benefits of fish attraction to fish populations and ecosystem services like fisheries; if fish attraction without increasing actual fish production occurs that only serves to aggregate fish making them easier catch (Eklöf et al., 2006; Hasselström et al., 2018; Corrigan et al., 2024b).

The aim of this study was to evaluate the ecological role of coastal kelp aquaculture as an artificial habitat during the pre-harvest period in Norway. Therefore our objectives were to: (1) assess fauna diversity and community composition associated using both eDNA and traditional sampling techniques; (2) compare biodiversity metrics between kelp farm sites and nearby natural habitats to determine the extent of habitat provisioning; and (3) understand how epifauna diversity varies between farm substrates given epifauna differences with natural areas have been previously well described (Bekkby et al., 2023). This information is vital to provide recommendations for sustainable aquaculture practices and to inform on the challenges and opportunities of biodiversity monitoring methods and their relevance for nature reporting in this emerging industry in Norway.

## METHODS

### 2.1 Field Sites

Field work was conducted during 5-18 April 2024. Two kelp farms along the Norwegian west coast were studied i.e. the ‘Trollsøy’ site near to Austevoll (Bergen) and ‘Inntian’ site off Frøya (Trondheim, Figure 1). Both locations cultivated S. latissima and A. esculenta at depths of ca. 2 m, over bottom depths of 7-30 m and harvested ca. 30-40,000 tonnes in the year of sampling (Figure 1C). Two habitat types were studied as control sites; these were one mixed S.latissima/L. hyperborea ‘kelp forest’ (ca. 5-10 m bottom depth), and one ‘pelagic’ control (approx. 30m bottom depth). Given that exposure is an important determinator of species abundance and diversity, sites were selected such that the exposure was as similar as possible between all three habitat types; this resulted in some control sites being closer to land than the kelp farm in the Austevoll location.

**Figure 1.**
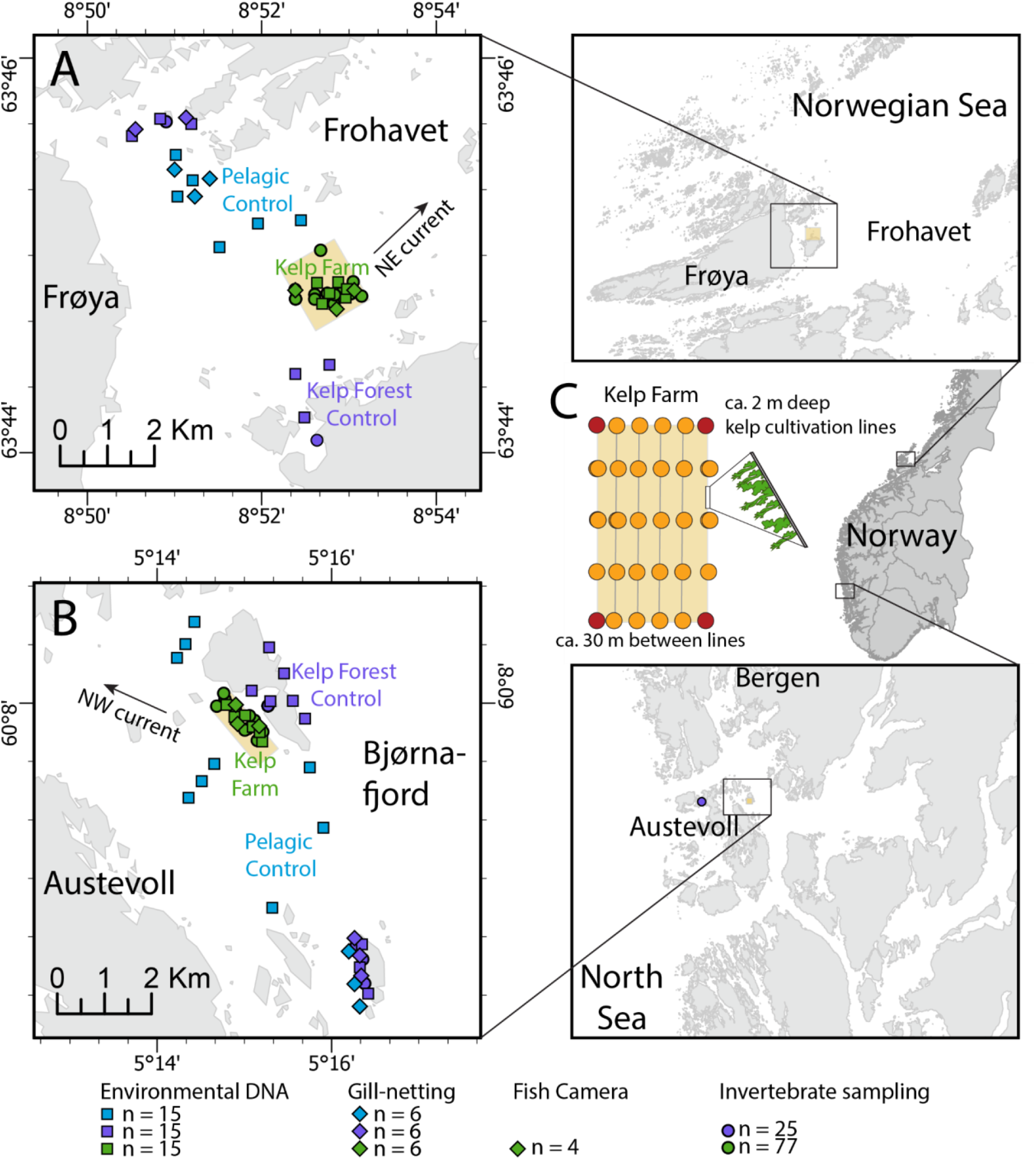
Map of sampling sites and habitat types at two Norwegian kelp aquaculture locations. A, B) Detailed site maps indicating the placement of kelp farms (cultivating *Saccharina latissima* and *Alaria esculenta*), natural kelp forest controls (*S. latissima/Laminaria hyperborea*), and pelagic control sites. Overview maps showing the locations of the two study sites: ‘Trollsøy’ near Austevoll (close to Bergen) and ‘Inntian’ near Frøya (close to Trondheim), along the Norwegian west coast. C) Schematic of the approximate kelp farm design, illustrating the relative depths of kelp cultivation and distance between lines.

### 2.2 Fishing and fish observations

Fishing was conducted with multi-mesh gill-nets (NORDIC gill-net) that were 30 m long and 1.5 m deep; divided in 2.5 m panels with mesh sizes 43.0, 19.5, 6.5, 10.0, 55.0, 8.0, 12.5, 24.0, 15.5, 5.0, 35.0, and 29.0 mm. Three nets were placed horizontally at a depth of 2–4 m in all three habitats; kelp farm, kelp forest control and pelagic control. Fishing occurred over 24 h for four days and nights in both Austevoll and Frøya (Table S1). Fish were collected early morning, identified to species and total length (TL) and weight measurements were recorded for each individual. All fish representing different nets in the same habitat type and mesh sizes were lumped and called one sample catch. In addition, fish presence on kelp during harvest was noted with a subset being hand-collected and preserved at −20°C to measure TL at a later date. In addition, two cameras were deployed at both kelp farms to capture fish diversity and behaviour during daylight hours using GoPro Hero 10 Black cameras, recording a photo every 10 seconds (Table S4). Other fishing methods like trammel and trapping were tried unsuccessfully as detailed in the Appendix.

### 2.3 Invertebrate collection and identification

Epifauna were collected on three replicates of *S. latissima* blades and holdfasts and by scraping 30 cm of production rope, structural rope (approximately 2 m depth) and floating plastic buoys (approximately 1m depth) between 21 February to 17 April 2025 at Austevoll and 12 March to 19 April at Frøya. *S. latissima* blades and stipes were also collected in natural kelp forest habitats at both areas in April, as control. The kelps and rope scrapes were collected from the boat, placing samples into plastic bags and subsequently freezing at −20°C.

In addition, mobile invertebrates were sampled using fauna traps, constructed by using a 1-m-long sisal rope (with 8 mm diameter), untwisted into its three parts, and bundled together with a strip (as (Bekkby et al., 2023)). The traps have been designed and tested to capture an assembly of species representative of the kelp forest–associated mobile invertebrate community, including crustaceans, gastropods, polychaetes, and bivalves (Christie et al., 2007). Fauna traps were deployed at the same locations in the farm and the kelp forests where we collected the kelp replicates (i.e. 18 fauna traps in total, half at the farm half at the control site). The traps were also put out at approximately 1-2 m depth and were retrieved after 72 h. The traps in the kelp farm and the wild kelp forests were collected from the boat (each trap was put gently into a plastic zip bag), by gently lifting traps up to the surface to reduce mobile invertebrate loss (Figure 2). Sampled kelps and fauna traps were rinsed in freshwater, and captured fauna was retained in a 250 µm sieve (Christie et al., 2003; Bekkby et al., 2023) and preserved in 70% ethanol.

**Figure 2.**
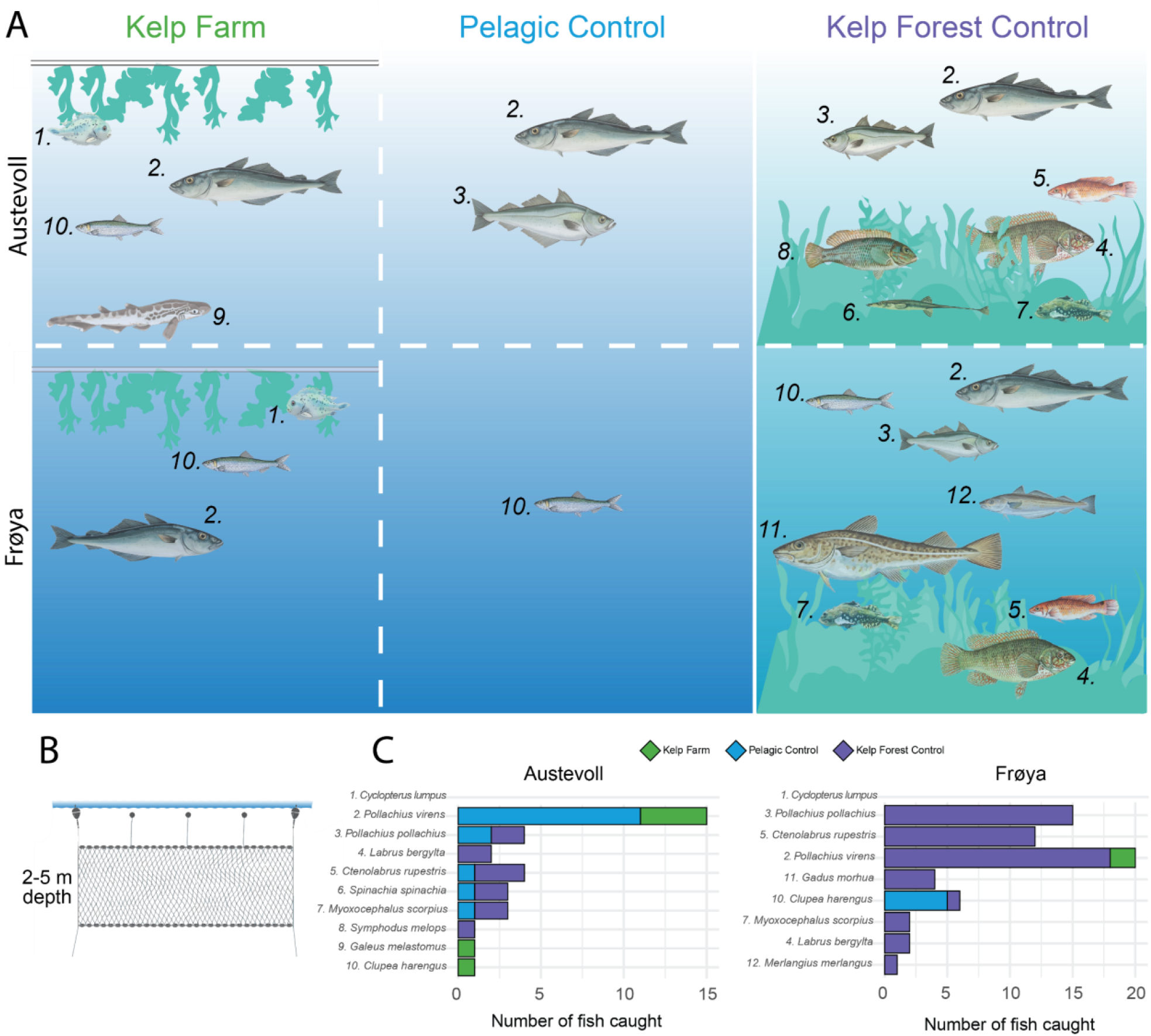
Traditionally observed fish community composition differences between the Kelp Farm and Control sites, surveyed using NORDIC gill-nets and hand collecting of 1. *Cyclopterus lumpus*. A) schematic of fish species diversity to compare Kelp Farm and Controls between Austevoll (top) sites and Frøya (bottom) sites, species (1-12) are indicated in C. B) simplified schematic of the positioning of 18 NORDIC gill-nets deployed over four nights, containing panels of varying mesh size. C) Barplots of counts of fish caught across sites, where counts hand collections of 1. *Cyclopterus lumpus*, the lumpfish are not shown given they were only collected at Kelp Farm sites *ad-hoc*.

The species were identified to the lowest possible taxonomic level using a Leica TL5000 Ergo Transmitted Light Base magnifier, based on the identification literature provided by Haywards and Ryland (1996) and sources further described in (Bekkby et al., 2023). The nomenclature used in the species list is in accordance with the database ‘Worlds Register of Marine Species’. The number of fauna taxa and individuals were counted and categorized into ranges if this exceeded 20 individuals of each species, then the median value was selected for analyses. Only taxa identified to family level were retained for statistical analyses with the exception of taxa identified as Nematoda and to sub-class level.

### 2.4 eDNA collection via water filtering

To obtain eDNA, triplicates of seawater samples (1.2 l) were collected at each habitat type (kelp farm, kelp forest control and pelagic control) from 2 m depth using Niskin bottles during two events (16, 18 April 2024) at Austevoll and three events (6, 8, 10 April 2024) at Frøya (Figure 1, Appendix Table S2). To better capture spatiotemporal biodiversity, seawater samples were taken at varying tidal heights, and locations between sampling events, upstream of kelp farms. All sampling equipment was sterilized with 10% bleach before each sampling event and thoroughly rinsed with seawater from the sampling point area before each use. All samples were collected and processed while wearing newly donned protective equipment such as nitrile gloves to prevent risk of contamination between samples or from outside sources. Seawater samples were maintained cool (4 °C) with no direct exposure to sunlight and were filtered within 5 hours of collection.

Seawater was filtered on site through 0.22 μm Sterivex filter units (Merck KGaA, Darmstadt, Germany) using a peristaltic pump (ST HandyPump, 300 rpm, DMR Miljø og Genoteknikk, Oslo, Norway). Pump hosing was cleaned by passing 10% bleach followed by 100 ml MilliQ Ultrapure water and then 200 ml of sample seawater, before beginning filtration of sample through the Sterivex unit. A total of 1 l of water was filtered through each of the triplicate filters from each location to ensure a standard volume between samples. After drying the filters by pumping air through using a sterile 50/60 ml syringe from BD Plastipak, the filters were placed in prelabeled sterile 50 ml Falcon tubes (Thermo Fisher Scientific, Waltham, MA, United States), and prelabeled bags for transport to UiT The Arctic University of Norway (UiT) and were kept at −80°C in an eDNA dedicated freezer. The syringes were changed, and operators’ hands were meticulously sterilized, between each sample using 10% bleach solution to limit contamination. A control blank was run on each sampling day to quantify contamination during the filtering process by filtering 0.5 l MilliQ Ultrapure water through a filter and drying the filter in the same manner as the samples.

### 2.5 eDNA laboratory analyses

The Sterivex filters used for water sampling underwent DNA extraction in over-pressured eDNA clean-labs at the Arctic University of Norway (UiT). In house protocols were followed that relied on vigilant care for cleanliness within and outside of the eDNA laboratories and avoidance of potential contaminant sources at UiT and personal life during the weeks of eDNA extraction lab use (Turon et al. 2022). Airborne DNA contamination risks were mitigated through use of a pressure positive eDNA extraction rooms and airlocked changing and sample preparation rooms. eDNA extraction protocols were meticulously followed for the modified use of DNEasy Blood and Tissue® (Qiagen, Hilden, Germany) kits. In short, a total of 500 μl of lysis buffer were added to each Sterivex filter, sealed with sterile caps at both ends, and incubated 24 h on a rotary wheel placed in a 56°C incubator oven to ensure full lysis of the particulates captured within the filter membrane. The lysed solution was then centrifuged out of the filter casing and into 2 ml Eppendorf tubes and the rest of the extraction followed the standard protocol. Subsequently each sample was eluted in 75 μl elution buffer, of which 20 μl was aliquoted for PCR, library preparation and sequencing.

A multiplexing approach was used for sequencing the 45 samples on an Illumina MiSeq next-generation sequencer (Illumina, San Diego, CA, United States). Amplicons were COI, and 12S. The partial COI Leray-XT fragment (313 bp) was amplified using the mlCOIintF-XT/jgHCO2198 primer pair (Wangensteen et al., 2018), while the 12S rRNA fragment (163-185 bp) was amplified using the MiFish-U primer pair (Miya et al., 2015). Samples included three PCR blanks as well as field blanks for each sampling event. 8-base tags were used to uniquely label each sample as in Wangensteen et al. (2018). PCR amplifications were conducted in 20 μl reactions containing 2 μl (COI), 3μl (12S), of DNA template, 10 μl of AmpliTaq Gold Master mix, 0.16 μl of Bovine Serum Albumin (20 μg/μl), 1 μl of each forward and reverse primer (5 μM) and 5.84 μl (COI), 4.84 μl (12S), of H2O. The PCR profiles were as follows, CO1: 95°C for 10 min; 35 cycles × (94°C/1 min, 45°C/1 min, 72°C/1 min); 72°C/5 min, 12S: 95°C for 10 min; 40 cycles × (95°C/ 30 sec, 60°C/ 30 sec, 72°C/ 30 sec); 72°C/5 min. Only one PCR replicate was run per sample, for COI, but 3 replicates were made for the 12S marker. The success of PCR amplifications was checked by gel electrophoresis in 1% agarose and PCR products were then pooled together into a multiplex sample pool for each amplicon. MinElute PCR purification columns (Qiagen) were used to concentrate the pooled DNA and to remove fragments below 70 bp. Library preparation was performed with the QIAseq 1-step amplicon library kit (Qiagen) with an adapted bead cleanup,and the exact library concentration was measured in a qPCR machine (ThermoFisher), using the NEBNext Library Quant Kit (New England BioLabs). Finally, pools were sequenced along with 1% PhiX on an Illumina Novaseq 6000 platform at Novogene (Cambridge, UK).

### 2.6 eDNA bioinformatic analyses

The Mjolnir v.1.2 pipeline (available from https://github.com/uit-metabarcoding/MJOLNIR) was used for bioinformatic processing of raw data. In short, the pipeline directed the following analyses for each primer set separately: The OBITools v1.01.22 software suite (Boyer et al., 2016) was used for the initial steps of the bioinformatic analyses. Paired-end reads were aligned using illumina paired-end and only sequences with alignment quality score > 40 were kept. Demultiplexing was done using the sample tags with ngsfilter, which also removed primer sequences. Aligned reads with length of 299–320 bp (CO1), and 140-190 bp (12S) and without ambiguous positions were selected using obigrep and then dereplicated with obiuniq. Chimeric sequences were removed using the uchime-denovo algorithm implemented in vsearch v1.10.1 (Rognes et al., 2016). Clustering of sequences into molecular operational taxonomic units (MOTUs) was performed using SWARM 2.0 (Mahé et al., 2015) with a d-value of 13 for CO1 (Bakker et al., 2019) and d-value of 1 for 12S following Mjolnir guidelines. Taxonomic assignment of the representative sequence of each MOTU was done with the ecotag algorithm (Boyer et al., 2016) using a local database of Leray or 12S fragment sequences (available from https://github.com/uit-metabarcoding/DUFA). Sequences for the MOTUs of interest (abundances > 0.5% of the total reads) were manually checked for a better match by BLAST search against the NCBI GenBank database, and best IDs were changed to reflect a higher percent match if one was found. Sequences assigned to contamination of terrestrial origin, and MOTUs that were present in the control samples with more than 10% of their total read abundance were removed. MOTUs with the same taxonomic assignment that resulted from intraspecific variation were merged. Prior to statistical analyses, non-bony fishes were removed from the final 12S list along with MOTUs of taxa not identified to genus or species level while non-metazoans were removed from the CO1 list along with MOTUs of taxa not identified to family, genus or species level.

### 2.7 Statistical analyses

Statistical analyses were performed in R version 3.1.3 (https://www.R-project.org/) with the vegan package [version 2.5–6; (Oksanen et al., 2019)] and graphic visualisations were done with ggplot2 package (Wickham, 2016).

#### 2.7.1 of traditionally collected fish and invertebrate data

No statistical analyses were performed on fishing or fish camera data given these data types were semi-qualitative. Instead, barplots of observation counts were generated for visual interpretation. Approximate (due to averages of ranges used for some species) abundances of traditionally sampled benthic invertebrate families per sample type and site were first analysed using non-metric multidimensional scaling (nMDS) with Bray-Curtis dissimilarities was performed with the metaMDS function in Vegan (Dixon, 2003) with 100 iterations.

Pairwise differences between sample type were tested using Wilcoxon rank-sum tests with Benjamini-Hochberg (BH) correction for multiple comparisons (Benjamini and Hochberg, 1995). Significant differences were annotated on the plots using the ggpubr package. Operator and Treemaps from treemapify package (Wilkins, 2021) were created to visualize the three most abundant invertebrate families per sample type and location.

#### 2.7.2 of eDNA data

eDNA sequence reads were first transformed to relative abundances to build a Bray-Curtis dissimilarity matrix, which was used to assess the variance in community composition using Permutational Multivariate Analyses of Variance (PERMANOVA). Samples were categorized as a function of Site (Kelp Forest Control, Kelp Farm, Pelagic Control), and Operator (Ocean Forest, Seaweed Solutions) and the univariate effects of these factors on the community composition were tested using adonis function with 999 permutations. Additionally, PERMDISP analysis (betadisper function) was performed for significant factors to determine if their effect was due to different multivariate mean or to different heterogeneity of the groups. Non-metric multidimensional scaling (nMDS) representation with Bray-Curtis dissimilarities was performed with the metaMDS function with 500 iterations. Shannon diversity and MOTU richness per sample were calculated in vegan (Dixon, 2003) after rarefaction to the lowest total number of reads per sample, to account for differences in sample sequencing depth. Then, two-way analysis of variance (ANOVA) was performed to detect significant differences between Date and Type in alpha diversity values.

An indicator species analysis (Dufrêne and Legendre, 1997) was performed in R using the labdsv package (Roberts, 2016) to detect potential associations of certain eukaryotic phyla to each type of site (Kelp Farm or Controls) per location (Austevoll or Frøya) and dataset (12S or CO1). Those with Indval values > 0.5 (p-value < 0.05 in all cases) were identified as indicator phyla. The top six (for 12S) and four (for CO1) MOTU Species’ and genus’ that differed in sequence read abundance between sites (Kelp Farm and Controls) were identified and pairwise differences between sites were tested using Wilcoxon rank-sum tests. Numbers differed in accordance between variation for each dataset. For the CO1 dataset, these taxa were restricted to fauna with benthic adult life stages to test for the influence of fouling the Kelp Farm structures. Benthic or pelagic (or bentho-pelagic for 12S data) adult life stage was identified for all taxa using online databases (fishbase.se, marinespecies.org). Treemaps from treemapify package (Wilkins, 2021) were created to visualize the overall eukaryotic composition of the sampling location considering read abundance and MOTU richness.

## 3. RESULTS

### 3.1 Fish Community Composition

Traditional sampling using NORDIC gill-nets and hand collection revealed distinct differences in fish community composition between kelp farm and control sites (Figure 2). Across both Austevoll and Frøya, fish diversity and abundance were generally lower at kelp farm sites where communities appeared pelagic. Notably, the lumpfish (juveniles, Cyclopterus lumpus, Appendix Table S3) was the only species observed exclusively at kelp farms, due to no survey effort at control sites. Fish were more frequently captured in control habitats, particularly in the kelp forest controls where typical benthic and benthic-pelagic fish were observed (e.g. Labrids, Myxocephalus, *Gadus morhua*), in addition to the pelagic component present at all sites (like saithe, *Pollachius virens* and herring, *Clupea harengus*, Supplementary File 1). Camera deployments at the kelp farms recorded a total of 13,936 images, with multiple observations of saithe at both sites. These fish were observed in small to moderate schools (1–32 individuals, Figure S1) and were not residentary nor observed feeding kelp farm associated invertebrates (Table S5). Despite the high number of images, fish diversity remained low, and no additional species were consistently observed across sampling events.

### 3.2 eDNA Community Composition

We analysed 69.8 million 12S and 24.7 million CO1 eDNA sequence reads, resulting in 165 and 3,655 MOTUs respectively after filtering. No MOTUs were identified as indicator species for kelp farms at either location. Only lemon sole (Microstomus kitt) was identified as an indicator species for the kelp forest control at Frøya (IndVal = 0.772, p = 0.041).

#### 12S Dataset

Non-metric multidimensional scaling (nMDS) of 12S data revealed moderate separation between sites, with kelp farm communities positioned between kelp forest and pelagic controls (Figure 3A). PERMANOVA indicated significant differences in community composition by site and operator (F = 6.9995, R^2^ = 0.344, p = 0.001). However, no individual MOTUs were significantly more abundant at kelp farms (Figure 3B). Alpha diversity analyses showed significant differences in Shannon diversity and MOTU richness between operators (p < 0.001), but not between sites (Figure 3C–D).

**Figure 3.**
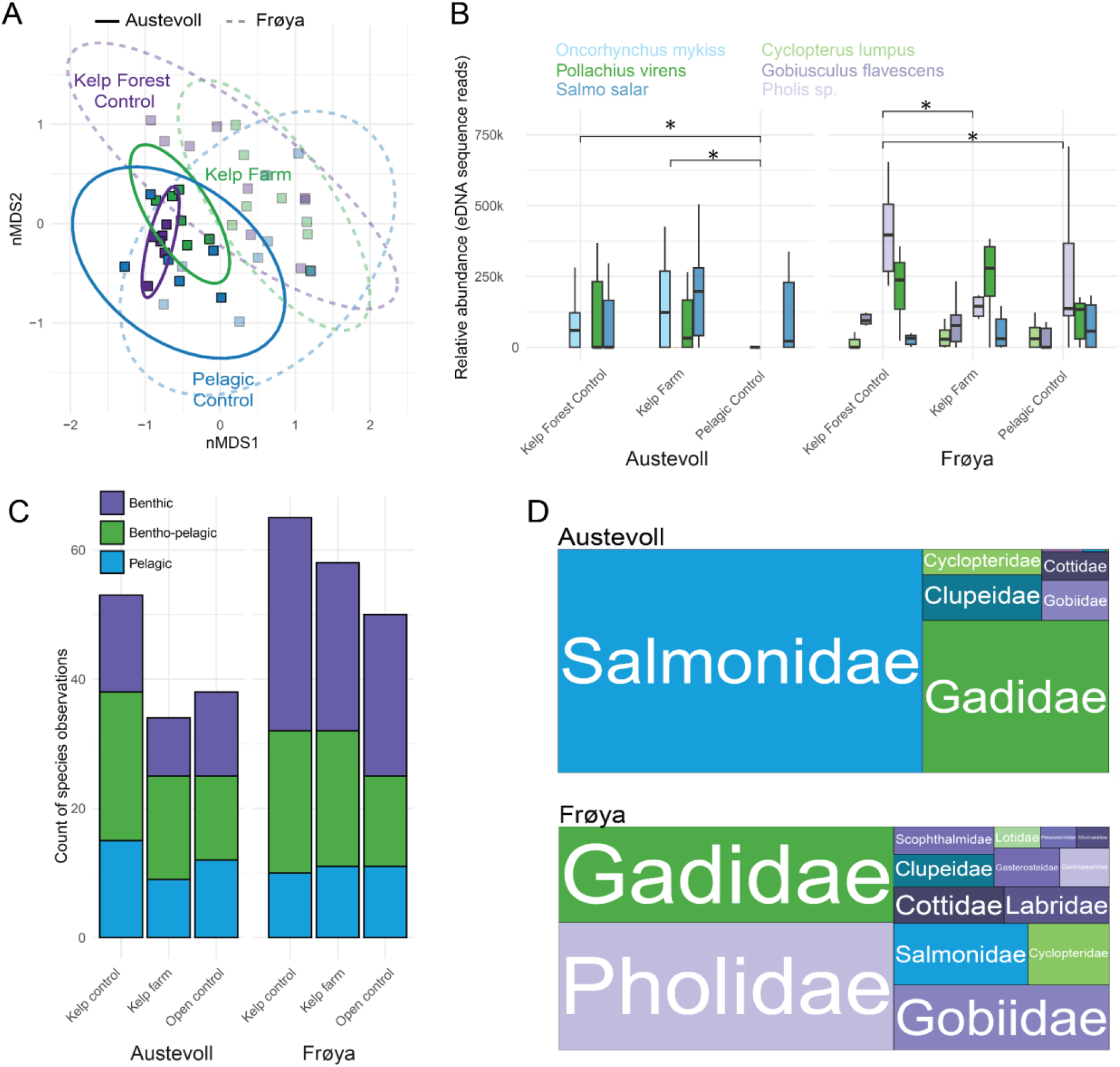
Environmental DNA (eDNA) 12S sequence diversity and relative abundance between the Kelp Farm and Control sites. A) nMDS of locations and sites (stress = 0.17, Austevoll, full; Frøya, semi-transparent and dashed). Ellipses display 95% confidence intervals (CI) of distributions. B) Relative abundance of sequence reads for select MOTU species showing differences between sites indicated by statistical significance (p>0.05: no asterix, p<0.05*, p<0.01**, p<0.001***). C) Barplots of the counts of benthic (purple), bentho-pelagic (green) and pelagic (blue) individuals observed at each site that show no significant differences at either Austevoll (left) or Frøya (right). D) treemap visualisation of the species diversity and abundance that scales with the sizes of 12S MOTUs overall at Austevoll (top) and Frøya (bottom). Colours reflect benthic vs. pelagic ecologies of MOTUs.

#### CO1 Dataset

The CO1 dataset, representing primarily invertebrate taxa, showed similar patterns. nMDS ordination indicated some separation by site and operator (Figure 4A), with PERMANOVA confirming significant effects (F = 2.6115, R^2^ = 0.160, p = 0.004). As with 12S, no MOTUs were significantly associated with kelp farms and of the benthic species showing the largest sequence read abundance differences between kelp farm and controls sites, none were statistically significant (Figure 4B). Alpha diversity analyses again showed significant differences by operator (Shannon: p = 0.0003; Richness: p = 3.47e-05), and a significant interaction between site and operator (Figure 4C–D). MOTUs were dominated by Archaeobalanidae (barnacles) and pelagic calanus species (Figure 4D).

**Figure 4.**
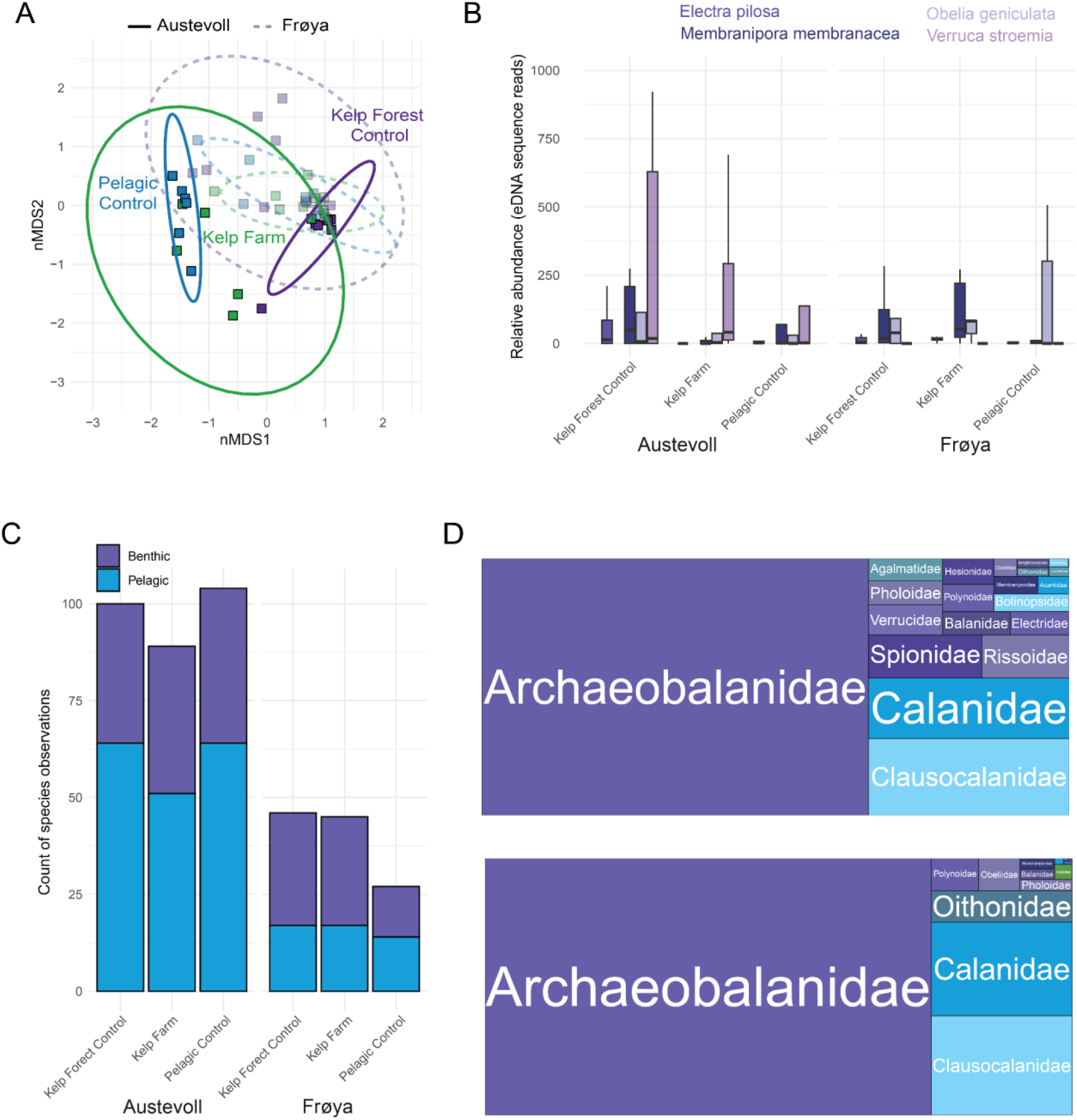
Environmental DNA (eDNA) CO1 sequence diversity and relative abundance between the Kelp Farm and Control sites. A) nMDS of locations and sites (stress = 0.12, Austevoll, full; Frøya, semi-transparent and dashed). Ellipses display 95% confidence intervals (CI) of distributions. B) Relative abundance of sequence reads for select MOTU species showing differences between sites indicated by statistical significance (p>0.05: no asterix, p<0.05*, p<0.01**, p<0.001***). C) Barplots of the counts of benthic (purple), and pelagic (blue) individuals observed at each site that show no significant differences at either Austevoll (left) or Frøya (right). D) treemap visualisation of the species diversity and abundance that scales with the sizes of CO1 MOTUs overall at Austevoll (top) and Frøya (bottom). Colours reflect benthic vs. pelagic ecologies of MOTUs.

### 3.3 Invertebrate Community Composition from Traditional Sampling

Traditional sampling of benthic invertebrates revealed significant differences in community composition between sample types (Figure 5A). nMDS ordination showed clear separation between farm-associated substrates (kelp, ropes, buoys) and natural kelp forest controls. Boxplots confirmed significantly lower richness and abundance in farmed kelp samples compared to wild kelp (Figure 5B). Treemaps illustrated that farmed kelp and associated structures supported fewer and less diverse invertebrate families than ropes and buoys that supported intertidal bivalves (Mytilidae) while Fauna Trap samples were dominated by copepods (Supplementary File 1, Figure 5C). Only one alien species, the Japanese ghost shrimp (Caprella mutica), was detected, only at kelp farm sites (Supplementary File 1) and only through traditional sampling.

**Figure 5.**
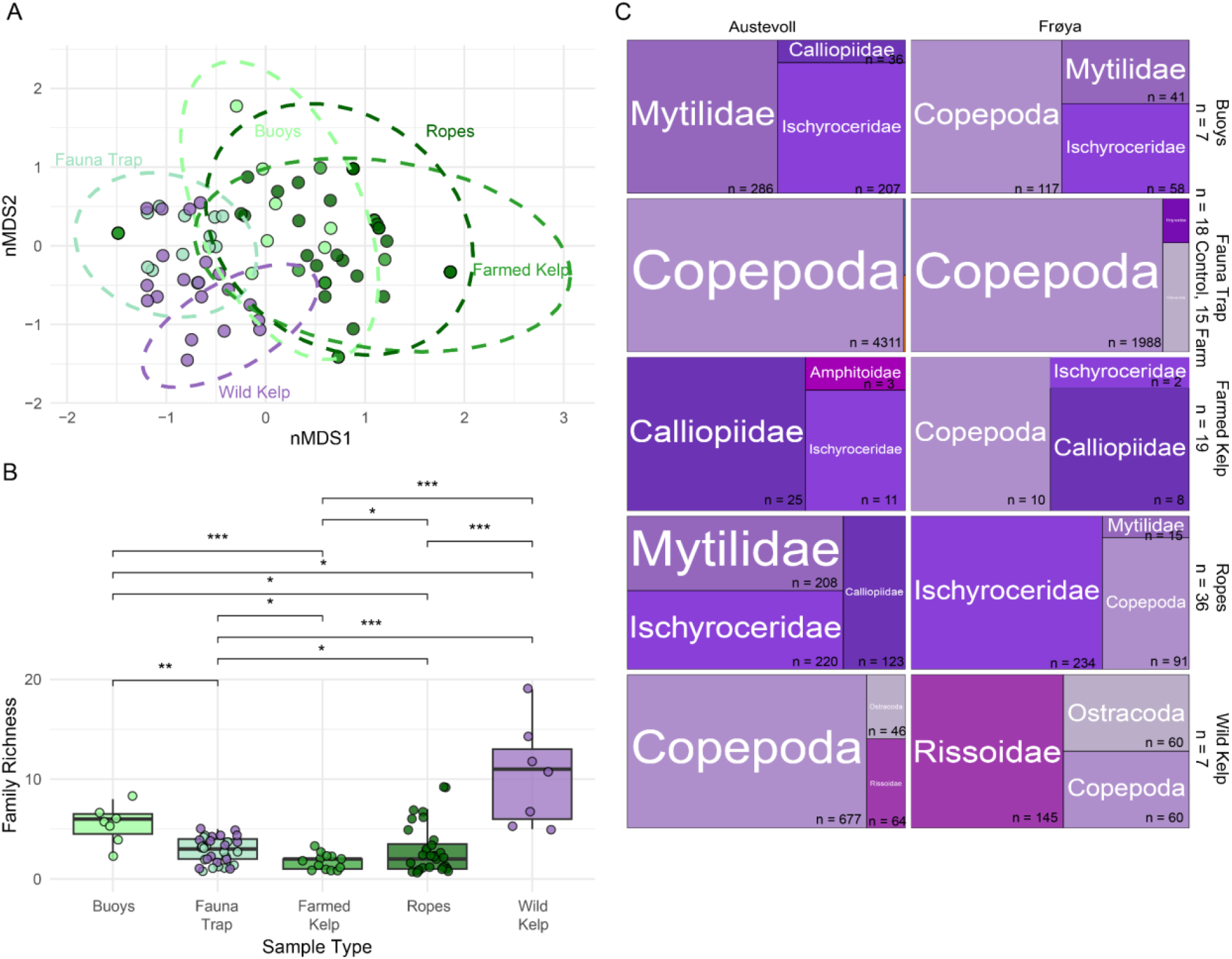
Traditionally sampled benthic invertebrate community composition differences across both locations between Sample Type (Buoys, Ropes, Fauna Trap, Farmed and Wild Kelp). A) nMDS of Sample Type (greens: kelp farm samples, purple: control samples, stress = 0.11). Ellipses display 95% confidence intervals (CI) of distributions. B) Boxplots of Sample type differences with statistical significance (p>0.05: no asterix, p<0.05*, p<0.01**, p<0.001***) and C) treemaps depicting the abundance of invertebrate families for each sample type between both locations showing the number (n) of individuals within each family.

## DISCUSSION

Our findings support the growing body of evidence that kelp aquaculture, in the Northeast Atlantic, provides limited habitat provisioning compared to natural kelp forests. Here we discuss factors driving these observations like current harvesting and operation regimes, in exposed waters and the implications of our results for management in terms of improving monitoring practices and nature reporting that are becoming increasingly relevant as the seaweed industry expands in the Atlantic (Duarte et al., 2021; Pardo et al., 2025).

### 4.1 Kelp farms as limited habitat providers

Through traditional sampling, we find that fish species diversity and abundance is lower at kelp farm sites compared to natural kelp forest control sites, instead it is more similar to pelagic control sites. The lumpfish was the only fish species present at kelp farms that exhibits a shallow-dwelling benthic behaviour that could theoretically benefit from the shelter and food provision of kelp aquaculture. This aligns with previous findings that kelp farms may not replicate the ecological complexity of natural blue forest habitats (Visch et al., 2020a; Christie et al., 2022).

Correspondingly, we found limited differences between kelp farm and control sites through eDNA sampling. 12S sequence MOTUs reveal no species as an indicator for the kelp farm or in significantly greater abundance there. Instead, sequence read abundance suggests eDNA at kelp farms comprises a mid-point between the near inshore kelp forest and pelagic controls. The same can be seen looking at individual species where the 12S eDNA read abundance of benthic species like the two-spotted goby (*Gobius flavescens*) is more similar between the kelp farm and kelp forest than the pelagic control while pelagically dispersing species like the lumpsucker show more similar read abundance between kelp farm and pelagic control sites. This agrees with (Schutt et al., 2023) who also reported minimal differences between kelp farms and control sites using eDNA.

We theorise limited fish attraction is driven by limited prey items i.e. our finding of significantly lower invertebrate richness in kelp farm samples than wild kelp samples, consistent with (Bekkby et al., 2023) and (Corrigan et al., 2024a). This was especially evident for the comparison with farmed kelp as can be expected due to relatively early harvesting times (May) currently employed to ensure a low-biofouled end product.

Invertebrate richness was lower but to a lesser degree in supporting ropes and buoys that had remained over several years. Literature points to these communities being likely driven by local species diversity that increases over time but is unlikely to represent those of natural kelp habitats even after prolonged periods of years (Eklöf et al., 2005). The lack of indicator species for kelp farms using CO1 eDNA and the absence of significant differences in CO1 alpha diversity between sites further reinforce the limited habitat provision. Key differences exist in invertebrate species between natural and farm environments e.g. the lack of Rissoidae and Lacuna snails in the farm (Figure 4C), that are a key component in the diet of wrasses (Labridae) and typically abundant in kelp forests in Norway (Christie et al., 2022), as also shown herein. These results are consistent with the idea that kelp farms, especially when harvested early to avoid fouling, may not provide sufficient time or complexity for kelp forest-associated communities to establish (Corrigan et al., 2024a).

In a Norwegian kelp forest, (Christie et al., 1998) found that it took more than 5 years for the species community associated with kelp to return after trawling, indicating the time needed for a kelp farm to function as a true kelp habitat. In the wild, the kelp-associated fauna increases in density and mobility during the warmer seasons, inhabiting a more heterogenous habitat with several macroalgae species (Norderhaug et al., 2002; Christie et al., 2003). Our results indicate that similar effects in kelp farms cannot be expected with an early harvesting time. Supporting this, (Corrigan et al., 2024b) observed higher fish abundance in kelp and mussel (Mytulis edulis) farms compared to control sites in September in the UK.

Farm size, wave exposure and other environmental conditions are factors influencing habitat provisioning (Bekkby et al. 2019; Broch et al. 2019; Gundersen et al. 2021) and will differ geographically, due to differences in industry maturity, local regulations and climates. In our study, both sites were relatively small scale, harvesting ca. 30-40 tonnes kelp in the year of sampling and were operating in relatively exposed locations. We cannot exclude the possibility that our results may have been different at larger scale farms. Biofouling on sugar kelp is known to be impacted by wave exposure (Visch et al., 2020b), where it would be higher in more sheltered environments and higher on artificial substrates not seeded by kelp for cultivation (Walls et al., 2019). However, it should be recognised that exposure is challenging to measure effects of between studies given that it is was not measured herein and as (Norderhaug et al. 2014) show, the richness of invertebrate communities decreases with exposure when exposure is low. Therefore we find it likely that more sheltered (but not low exposure) kelp farms in Norway, operated closer to the coast, may present increases in the degree of habitat provision for invertebrates and fish compared to the studied sites herein. The presence of Caprella mutica, an alien amphipod species, at kelp farms further supports the idea that artificial substrates can facilitate alien (non-native) species colonization, likely due to the reduced competition that provides opportunities compared to challenges of colonizing climax (mature) wild habitats (Ashton et al., 2007).

### 4.2 Limitations and improvements for monitoring

Despite our sampling approach being limited in duration, our observations are unlikely to have been heavily biased given their agreement with Grimm Torstensen (2020) who show similar fish biodiversity and abundance at kelp forest and pelagic control sites in April and September at Frøya. The fishing effort placed in the kelp farms was higher than control sites where in addition to gill nets, trammel nets and traps were also placed (see Appendix X). In addition we repeated fish hand-collection at harvest and camera operations in May 2025 at the Austevoll site with the similar result i.e. saithe and lumpfish being the only species recovered in similar numbers as in 2024 (Appendix Figure S2, Table S6). Fewer fish at kelp farms in the current compared to other studies could be explained partly by sampling choices e.g. compared to Corrigan et al., 2024b, who fished throughout the water column with hook and line and attracted fish to the kelp farm using baited cameras, we used passive methods that we suggest could more accurately reflect the impact of farmed biomass suspended in the water column that could in theory act as shelter or food provision.

eDNA did not find differences in communities that were detected by more traditional sampling methods. This is most likely because the traditional methods sample, for certain, organisms in the different habitats and structures, while eDNA sample in the water masses, most potentially tracing species not within the habitat and structures but transported from some distance away through the water masses. Our eDNA approach could be improved to increase sequence read abundance, avoid over-representation of some groups and increase the number of samples (Figure 3B, 4B). Increased sampling effort could be employed to better capture spatiotemporal eDNA dynamics. Overall, there is a large challenge performing eDNA monitoring at exposed kelp farms in Norway considering their high dynamics and variable current directions. Again, higher sampling resolution would likely aid the capture of local communities in addition to passive sampler methods to collect DNA over longer time periods, and greater sequencing depth. Greater sampling depth is especially required to overcome the large abundance of salmon (Salmo salar) and rainbow trout (Oncorhynchus mykiss) sequence reads likely from aquaculture that oversaturate during sequencing. We theorise increased saithe read abundance is probably biased by their attraction at nearby salmon aquaculture pens as widely acknowledged and shown by Turon et al. (2022), but saithe is clearly also abundant as traditional sampling at control sites and farms indicates.

Improved and standard eDNA practices may contribute to improved coherence between traditional and molecular sampling that may be important to more automated low-cost monitoring required by the industry. As an example; while traditional sampling detected *Caprella mutica*, eDNA did not, highlighting a mismatch between methods. This discrepancy may stem from low DNA shedding rates of invertebrates or dynamic water conditions at exposed sites, which dilute eDNA signals. Similar challenges have been noted in other studies using eDNA to monitor biofouling and invasive species (Bernard et al., 2019; Turon et al., 2022). To tackle this, species-specific assays targeting key taxa like biofouling herbivore pests e.g. Lacuna vincta or unwanted alien species such as *Caprella mutica*, are recommended but then result in an ability to capture the entire community (Blaine et al., 2025). These approaches could however enhance detection of rare or low-abundance species and better inform ecological and economic risk assessments.

### 4.3 Management Considerations

Although current kelp farms may not significantly enhance biodiversity, this limited impact can be viewed positively in terms of minimizing ecological disruption. As noted by (WWF, 2025) a lack of effect may still align with sustainability goals, especially when compared to more intensive aquaculture practices. However, the presence of alien species requires action to define guidelines (like (Wilding et al. 2021)) that identify their impacts to limit their colonisation, spread that could be deemed as a nature positive action that also aids the achievement of international biodiversity targets under the Kunming–Montreal Global Biodiversity Framework or national ones e.g. Norway’s ‘Ocean Management Plan’ (Meld. St. (2023-2024); www.regjeringen.no).

Policy should be developed surrounding the impacts of fish attraction (Methratta and Dardick, 2019), taking into account whether food provision occurs. Artificial habitats and additional efforts to increase biodiversity through placement of artificial reefs may result in unwanted and negative impacts, such as ecological-traps that hinder ecosystem restoration (Reubens et al., 2013; Methratta and Dardick, 2019; Rouse et al., 2019; Pardo et al., 2025). Concerns for biodiversity arise given that impacts of fish attraction are poorly understood and that local population and ecosystem vulnerability is not represented by vulnerability frameworks that occur at the species level e.g. IUCN red list (Bennun et al., 2024). Similar to fish aggregation effects of offshore energy structures, the attraction of fishes is not negative or positive per se (Coates et al., 2016). It depends on the actual benefits to populations in biomass increases as a result of secondary production (e.g. prey) and support of natural behaviours (Hasselström et al., 2018), requiring detailed scientific study of population structure, local dynamics, natural niches and behaviours that exceeded the remit of this study before careful consideration by authorities on which effects are deemed to be positive or not. We show the only relevant fish species present in Norwegian kelp farms from a nature-positive perspective appears to be juvenile lumpfish given that pelagic species like saithe and herring do not exhibit stationary foraging.

We were unable herein to provide an estimation of how many lumpfish may be present in kelp farms but we note their presence appears ubiquitous in Nordic countries as expected from their ecology as a shallow benthic-dwelling species as juveniles that can disperse long distances to reach farms (Visch et al., 2020a). (Visch et al., 2020a) estimated approximately 3000 lumpfish per ha−1 estimated in a kelp farm off Sweden. Considering the dispersal ability of lumpfish to migrate to nearby natural habitats during and after harvesting (that is a key factor in the potential of kelp farms to impact fish populations (Corrigan et al., 2024b)), further research would reveal this potential nature positive contribution by assessing the densities of lumpfish present in Nordic farms, their ability to benefit from prey items at the kelp farm and the potential for secondary production for the population they pertain to.

### 4.4 Conclusion

This study provides the first empirical comparison of fish and invertebrate communities at kelp farms and natural habitats in Norway, showing that current kelp aquaculture offers limited habitat provisioning. While this may reduce ecological risks, it also suggests that kelp farms are not ecological substitutes for natural kelp forests. Greater recognition and support from relevant authorities could help realize the environmental potential of low-trophic aquaculture by determining its nature-positive contributions. As we show, kelp farms are not inherently nature positive or negative—potential benefits like increased biofouling and fish attraction that may be increased with later harvesting schedules than are currently employed must be weighed against economic and ecological risks such as the reduced crop quality, seabed impacts and alien species spread that would coincide with such schedules. Future research aims towards controls for alien species, develop environmental impact assessments to include nature positive or negative contributions locally in collaboration with authorities, develop monitoring to identify population-level effects for relevant species like the lumpfish rather than species presence and assess how operating practices like farm scale and harvesting timing impact ecological outcomes.

## AUTHOR CONTRIBUTIONS

AJA and TB designed the study. AJA, HC, ØT, RB and LL collected samples for analysis. AJA, HC, KP, MB and RGT conducted the laboratory work. AJA analysed the data. AJA wrote the manuscript. All authors revised the manuscript.

## ACKNOWLEDGEMENTS

This work was developed by NIVA as a contribution to the Innovasjon Norge and Norges Forskningsråd Grønn Platform project ‘New products from cultivated seaweed for blue-green value-chains’ (no. 340964). We thank Seaweed Solutions AS and Ocean Forest AS for aiding sample collection. We thank two UiO ‘Arbeids Praksis’ students, Emma Ulvær and Emil Lange for their laboratory assistance. A permit to fish in the Trollsøy lobster sanctuary was granted by Fiskeri Direktoratet (24/3122 of 26.03.2024).

## DATA AVAILABILITY

DNA sequence data has been uploaded to the European Nucleotide Archive and can be accessed at the accession PRJEB96877 on acceptance of the article.

