## Supplementary material for "Combining environmental DNA and traditional sampling to assess the role of kelp aquaculture as an artificial habitat in Norway": 5. Supplementary Appendix

**Table S1. Gillnet sampling details.** OF=Ocean Forest. SES= Seaweed Solutions.

| Name | Latitude | Longitude | Type | Bottom depth | Site | Operator | Date in | Time in | Date out | Time out |
|---|---|---|---|---|---|---|---|---|---|---|
| Block 1 | 60.13369 | 5.24629 | Kelp farm | 24 | Trollsøy 2 | OF | 05/04/2024 | 16:00 | 09/04/2024 | 17:00 |
| Block 3 | 60.13276 | 5.24827 | Kelp farm | 29 | Trollsøy 2 | OF | 05/04/2024 | 16:30 | 09/04/2024 | 17:30 |
| Block 19 | 60.13089 | 5.25068 | Kelp farm | 49 | Trollsøy 2 | OF | 05/04/2024 | 17:00 | 09/04/2024 | 18:00 |

| | | | | | | | | | | |
|---|---|---|---|---|---|---|---|---|---|---|
| Kelp control 1 | 60.10897 | 5.27262 | Kelp Control | 10-15 m | Trollsøy 2 | OF | 06/04/2024 | 15:00 | 09/04/2024 | 10:00 |
| Kelp control 2 | 60.10667 | 5.27299 | Kelp Control | 10-15 m | Trollsøy 2 | OF | 06/04/2024 | 15:30 | 09/04/2024 | 10:30 |
| Kelp control 3 | 60.11032 | 5.27139 | Kelp Control | 10-15 m | Trollsøy 2 | OF | 06/04/2024 | 16:00 | 09/04/2024 | 11:00 |
| Deep control 1 | 60.10994 | 5.27077 | Pelagic Control | 25-30 m | Trollsøy 2 | OF | 06/04/2024 | 19:00 | 09/04/2024 | 11:30 |
| Deep control 2 | 60.10705 | 5.27229 | Pelagic Control | 25-30 m | Trollsøy 2 | OF | 06/04/2024 | 18:00 | 09/04/2024 | 12:00 |
| Deep control 3 | 60.10460 | 5.27282 | Pelagic Control | 25-30 m | Trollsøy 2 | OF | 06/04/2024 | 18:30 | 09/04/2024 | 12:30 |
| block 9 north | 63.74669 | 8.87677 | Kelp farm | 30-40m | Inntian | SES | 13/04/2024 | 13:00 | 17/04/2024 | 11:00 |
| block 9 south | 63.74631, | 8.87676 | Kelp farm | 30-40m | Inntian | SES | 13/04/2024 | 13:30 | 17/04/2024 | 11:00 |
| block 5 south | 63.74680 | 8.87916 | Kelp farm | 30-40m | Inntian | SES | 13/04/2024 | 14:00 | 17/04/2024 | 11:30 |

| | | | | | | | | | | |
|---|---|---|---|---|---|---|---|---|---|---|
| Deep control 1 | 63.75994 | 5.84760 | Pelagic Control | 25-30m | Inntian | SES | 13/04/2024 | 16:30 | 17/04/2024 | 12:00 |
| Deep control 2 | 63.75704 | 8.85105 | Pelagic Control | 25-30m | Inntian | SES | 13/04/2024 | 17:00 | 17/04/2024 | 12:00 |
| Deep control 3 | 63.76078 | 8.84250 | Pelagic Control | 25-30m | Inntian | SES | 13/04/2024 | 17:30 | 17/04/2024 | 12:30 |
| Kelp control 1 | 63.16136 | 8.84739 | Kelp Control | 7-10m | Inntian | SES | 13/04/2024 | 18:00 | 17/04/2024 | 13:30 |
| Kelp control 2 | 63.76152 | 8.85202 | Kelp Control | 7-10m | Inntian | SES | 13/04/2024 | 18:30 | 17/04/2024 | 13:30 |
| Kelp control 3 | 63.75898 | 8.85172 | Kelp Control | 7-10m | Inntian | SES | 13/04/2024 | 18:30 | 17/04/2024 | 13:30 |
| kelp trammel 1 block 13 north | 63.74712 | 8.87302 | Kelp farm | 30-40m | Inntian | SES | 14/04/2024 | 12:00 | 17/04/2024 | 10:30 |
| kelp trammel 2 block 13 south | 63.746330 | 8.873116 | Kelp farm | 30-40m | Inntian | SES | 14/04/2024 | 12:00 | 17/04/2024 | 10:30 |

**Table S2. eDNA filter sample details.** OF=Ocean Forest. SES= Seaweed Solutions.

| Sample ID | Latitude | Longitude | Site | Operator | Date | Time | Low tide time | Depth | Number | ID |
|---|---|---|---|---|---|---|---|---|---|---|
| 1 | 60.13096 | 5.25278 | Trollsøy 2 | OF | 06/04/2024 | 11:00 | 18:00 | 18.3m | 1 | Kelp control |
| 2 | 60.10675 | 5.27207 | Trollsøy 2 | OF | 06/04/2024 | 11:00 | 18:00 | 16.2m | 2 | Kelp control |
| 3 | 60.10921 | 5.27236 | Trollsøy 2 | OF | 06/04/2024 | 11:00 | 18:00 | 6m | 3 | Kelp control |
| 4 | 60.11623 | 5.26630 | Trollsøy 2 | OF | 06/04/2024 | 11:30 | 18:00 | 142m | 4 | Open control |
| 5 | 60.12149 | 5.26413 | Trollsøy 2 | OF | 06/04/2024 | 11:30 | 18:00 | 220m | 5 | Open control |
| 6 | 60.12766 | 5.26202 | Trollsøy 2 | OF | 06/04/2024 | 11:30 | 18:00 | 23m | 6 | Open control |
| 7 | 60.13168 | 5.25064 | Trollsøy 2 | OF | 06/04/2024 | 12:00 | 18:00 | 38m | 7 | Kelp farm |
| 8 | 60.13319 | 5.24657 | Trollsøy 2 | OF | 06/04/2024 | 12:00 | 18:00 | 26m | 8 | Kelp farm |

| | | | | | | | | | | |
|---|---|---|---|---|---|---|---|---|---|---|
| 9 | 60.13300 | 5.24546 | Trollsøy 2 | OF | 06/04/2024 | 12:00 | 18:00 | 16.5m | 9 | Kelp farm |
| 10 | 60.11385 | 5.25514 | Trollsøy 2 | OF | 08/04/2024 | 17:00 | 18:30 | in middle of bay | 10 | Kelp control |
| 11 | 60.134443 | 5.251314 | Trollsøy 2 | OF | 08/04/2024 | 17:00 | 18:30 | left side of bay close to kelp | 11 | Kelp control |
| 12 | 60.13307 | 5.25453 | Trollsøy 2 | OF | 08/04/2024 | 17:00 | 18:30 | 6m | 12 | Kelp control |
| 13 | 60.133303 | 5.24413 | Trollsøy 2 | OF | 08/04/2024 | 17:00 | 18:30 | | 13 | Kelp farm |
| 14 | 60.131561 | 5.24823 | Trollsøy 2 | OF | 08/04/2024 | 17:00 | 18:30 | | 14 | Kelp farm |
| 15 | 60.12979 | 5.25242 | Trollsøy 2 | OF | 08/04/2024 | 17:00 | 18:30 | | 15 | Kelp farm |
| 16 | 60.12751 | 5.24483 | Trollsøy 2 | OF | 08/04/2024 | 18:00 | 18:30 | 48 | 16 | Open control |
| 17 | 60.12585 | 5.24249 | Trollsøy 2 | OF | 08/04/2024 | 18:00 | 18:30 | 156 | 17 | Open control |

| | | | | | | | | | | |
|---|---|---|---|---|---|---|---|---|---|---|
| 18 | 60.12429 | 5.23972 | Trollsøy 2 | OF | 08/04/2024 | 18:00 | 18:30 | | 18 | Open control |
| 19 | 60.13184 | 5.26157 | Trollsøy 2 | OF | 10/04/2024 | 18:00 | 18:30 | 14 | 19 | Kelp control |
| 20 | 60.13468 | 5.25888 | Trollsøy 2 | OF | 10/04/2024 | 17:30 | 18:30 | 6 | 20 | Kelp control |
| 21 | 60.13826 | 5.25583 | Trollsøy 2 | OF | 10/04/2024 | 17:30 | 18:30 | 10 | 21 | Kelp control |
| 22 | 60.14058 | 5.24160 | Trollsøy 2 | OF | 10/04/2024 | 17:30 | 18:30 | 100 | 22 | Open control |
| 23 | 60.13889 | 5.23979 | Trollsøy 2 | OF | 10/04/2024 | 18:00 | 18:30 | 74 | 23 | Open control |
| 24 | 60.13767 | 5.23884 | Trollsøy 2 | OF | 10/04/2024 | 18:00 | 18:30 | 92 | 24 | Open control |
| 25 | 60.13219 | 5.24869 | Trollsøy 2 | OF | 10/04/2024 | 18:30 | 18:30 | 19 | 25 | Kelp farm |
| 26 | 60.13133 | 5.25208 | Trollsøy 2 | OF | 10/04/2024 | 18:30 | 18:30 | 40 | 26 | Kelp farm |
| 27 | 60.12991 | 5.25453 | Trollsøy 2 | OF | 10/04/2024 | 18:30 | 18:30 | 42 | 27 | Kelp farm |

| | | | | | | | | | | |
|---|---|---|---|---|---|---|---|---|---|---|
| Contol | NA | NA | Trollsøy 2 | OF | 06/04/2024 | | | | Contol | |
| Contol | NA | NA | Trollsøy 2 | OF | 08/04/2024 | | | | Contol | |
| Contol | NA | NA | Trollsøy 2 | OF | 10/04/2024 | | | | Contol | |
| SES 1 | 63.76112 | 8.85259 | Inntian | SES | 16/04/2024 | 17:00 | 16:30 | 6 | 1 | Kelp control |
| SES 2 | 63.76129 | 8.84839 | Inntian | SES | 16/04/2024 | 17:00 | 16:30 | 7 | 2 | Kelp control |
| SES 3 | 63.75856 | 8.85246 | Inntian | SES | 16/04/2024 | 17:00 | 16:30 | 7 | 3 | Kelp control |
| SES 4 | 63.76055 | 8.84277 | Inntian | SES | 16/04/2024 | 17:30 | 16:30 | 30 | 4 | Open control |
| SES 5 | 63.75436 | 8.85358 | Inntian | SES | 16/04/2024 | 17:30 | 16:30 | 46m | 5 | Open control |
| SES 6 | 63.75071 | 8.85940 | Inntian | SES | 16/04/2024 | 17:30 | 16:30 | 25.5 | 6 | Open control |

| | | | | | | | | | | |
|---|---|---|---|---|---|---|---|---|---|---|
| SES 7 | 63.74778 | 8.88412 | Inntian | SES | 16/04/2024 | 18:00 | 16:30 | | 7 | Kelp farm |
| SES 8 | 63.74682 | 8.88181 | Inntian | SES | 16/04/2024 | 18:00 | 16:30 | | 8 | Kelp farm |
| SES 9 | 63.74591 | 8.88045 | Inntian | SES | 16/04/2024 | 18:00 | 16:30 | | 9 | Kelp farm |
| SES Contol | NA | NA | Inntian | SES | 16/04/2024 | | | | Contol | |
| SES Contol | NA | NA | Inntian | SES | 18/04/2024 | | | | Contol | |
| SES 10 | 63.73620 | 8.87581 | Inntian | SES | 18/04/2024 | 15:00 | 17:00 | 7 | 10 | Kelp control |
| SES 11 | 63.73971 | 8.87353 | Inntian | SES | 18/04/2024 | 15:00 | 17:00 | 4 | 11 | Kelp control |
| SES 12 | 63.74077 | 8.88011 | Inntian | SES | 18/04/2024 | 15:00 | 17:00 | 6 | 12 | Kelp control |
| SES 13 | 63.74657 | 8.88573 | Inntian | SES | 18/04/2024 | 15:00 | 17:00 | 15 | 13 | Kelp farm |
| SES 14 | 63.74678 | 8.88373 | Inntian | SES | 18/04/2024 | 15:00 | 17:00 | 25 | 14 | Kelp farm |
| SES 15 | 63.74668 | 8.88059 | Inntian | SES | 18/04/2024 | 15:00 | 17:00 | 28.5 | 15 | Kelp farm |
| SES 16 | 63.75044 | 8.87790 | Inntian | SES | 18/04/2024 | 15:00 | 17:00 | 27 | 16 | Open |

| | | | | | | | | | | |
|---|---|---|---|---|---|---|---|---|---|---|
| | | | | | | | | | | control |
| SES 17 | 63.75306 | 8.87427 | Inntian | SES | 18/04/2024 | 15:00 | 17:00 | 17 | 17 | Open control |
| SES 18 | 63.75288 | 8.86670 | Inntian | SES | 18/04/2024 | 15:00 | 17:00 | 37 | 18 | Open control |

**Lumpfish numbers and sizes**

Lumpfish were collected ad-hoc from Austevoll in 2024 by the operator but were not measured or retained. Approximate sizes of 20-30mm were present suggesting these were juvenile individuals. Approximately 25 were observed on farmed kelp as it was taken aboard the harvest vessel, during two days of harvesting.

Numbers of lumpfish were similarly not counted at Frøya but sampled ad-hoc as observed on kelp during harvest. In total this resulted in 55 lumpfish being collected over two sampling dates at the Inntian kelp farm site. Total length was measured and ranged from 21-95 mm (Table S3).

**Table S3. Hand-collections of fish at the Frøya kelp farm during harvest in May 2024.** SES= Seaweed Solutions-

| Date | Site | Operator | TL (mm) | ID | Species |
|---|---|---|---|---|---|
| 30/05/2024 | Inntian | SES | 35 | 1 | *Cyclopterus lumpus* |
| 30/05/2024 | Inntian | SES | 36 | 2 | *Cyclopterus lumpus* |

| | | | | | |
|---|---|---|---|---|---|
| 30/05/2024 | Inntian | SES | 37 | 3 | *Cyclopterus lumpus* |
| 30/05/2024 | Inntian | SES | 37 | 4 | *Cyclopterus lumpus* |
| 30/05/2024 | Inntian | SES | 30 | 5 | *Cyclopterus lumpus* |
| 30/05/2024 | Inntian | SES | 31 | 6 | *Cyclopterus lumpus* |
| 30/05/2024 | Inntian | SES | 33 | 7 | *Cyclopterus lumpus* |
| 30/05/2024 | Inntian | SES | 31 | 8 | *Cyclopterus lumpus* |
| 30/05/2024 | Inntian | SES | 29 | 9 | *Cyclopterus lumpus* |
| 30/05/2024 | Inntian | SES | 28 | 10 | *Cyclopterus lumpus* |
| 30/05/2024 | Inntian | SES | 29 | 11 | *Cyclopterus lumpus* |
| 30/05/2024 | Inntian | SES | 29 | 12 | *Cyclopterus lumpus* |
| 30/05/2024 | Inntian | SES | 31 | 13 | *Cyclopterus lumpus* |
| 30/05/2024 | Inntian | SES | 30 | 14 | *Cyclopterus lumpus* |
| 30/05/2024 | Inntian | SES | 26 | 15 | *Cyclopterus lumpus* |

| | | | | | |
|---|---|---|---|---|---|
| 30/05/2024 | Inntian | SES | 28 | 16 | *Cyclopterus lumpus* |
| 30/05/2024 | Inntian | SES | 30 | 17 | *Cyclopterus lumpus* |
| 30/05/2024 | Inntian | SES | 30 | 18 | *Cyclopterus lumpus* |
| 30/05/2024 | Inntian | SES | 26 | 19 | *Cyclopterus lumpus* |
| 30/05/2024 | Inntian | SES | 25 | 20 | *Cyclopterus lumpus* |
| 30/05/2024 | Inntian | SES | 30 | 21 | *Cyclopterus lumpus* |
| 30/05/2024 | Inntian | SES | 25 | 22 | *Cyclopterus lumpus* |
| 30/05/2024 | Inntian | SES | 29 | 23 | *Cyclopterus lumpus* |
| 30/05/2024 | Inntian | SES | 27 | 24 | *Cyclopterus lumpus* |
| 30/05/2024 | Inntian | SES | 29 | 25 | *Cyclopterus lumpus* |
| 30/05/2024 | Inntian | SES | 28 | 26 | *Cyclopterus lumpus* |
| 30/05/2024 | Inntian | SES | 28 | 27 | *Cyclopterus lumpus* |
| 30/05/2024 | Inntian | SES | 27 | 28 | *Cyclopterus lumpus* |

| | | | | | |
|---|---|---|---|---|---|
| 30/05/2024 | Inntian | SES | 27 | 29 | *Cyclopterus lumpus* |
| 30/05/2024 | Inntian | SES | 26 | 30 | *Cyclopterus lumpus* |
| 30/05/2024 | Inntian | SES | 26 | 31 | *Cyclopterus lumpus* |
| 30/05/2024 | Inntian | SES | 22 | 32 | *Cyclopterus lumpus* |
| 30/05/2024 | Inntian | SES | 21 | 33 | *Cyclopterus lumpus* |
| 30/05/2024 | Inntian | SES | 27 | 34 | *Cyclopterus lumpus* |
| 30/05/2024 | Inntian | SES | 23 | 35 | *Cyclopterus lumpus* |
| 30/05/2024 | Inntian | SES | 24 | 36 | *Cyclopterus lumpus* |
| 30/05/2024 | Inntian | SES | 21 | 37 | *Cyclopterus lumpus* |
| 30/05/2024 | Inntian | SES | 22 | 38 | *Cyclopterus lumpus* |
| 30/05/2024 | Inntian | SES | 19 | 39 | *Cyclopterus lumpus* |
| 05/06/2024 | Inntian | SES | 21 | 40 | *Cyclopterus lumpus* |
| 05/06/2024 | Inntian | SES | 24 | 41 | *Cyclopterus lumpus* |

| | | | | | |
|---|---|---|---|---|---|
| 05/06/2024 | Inntian | SES | 31 | 42 | *Cyclopterus lumpus* |
| 29/05/2024 | Inntian | SES | 24 | 43 | *Cyclopterus lumpus* |
| 29/05/2024 | Inntian | SES | 25 | 44 | *Cyclopterus lumpus* |
| 29/05/2024 | Inntian | SES | 24 | 45 | *Cyclopterus lumpus* |
| 29/05/2024 | Inntian | SES | 26 | 46 | *Cyclopterus lumpus* |
| 29/05/2024 | Inntian | SES | 26 | 47 | *Cyclopterus lumpus* |
| 29/05/2024 | Inntian | SES | 32 | 48 | *Cyclopterus lumpus* |
| 29/05/2024 | Inntian | SES | 31 | 49 | *Cyclopterus lumpus* |
| 29/05/2024 | Inntian | SES | 37 | 50 | *Cyclopterus lumpus* |
| 29/05/2024 | Inntian | SES | 37 | 51 | *Cyclopterus lumpus* |
| 29/05/2024 | Inntian | SES | 39 | 52 | *Cyclopterus lumpus* |
| 29/05/2024 | Inntian | SES | 40 | 53 | *Cyclopterus lumpus* |
| 29/05/2024 | Inntian | SES | 40 | 54 | *Cyclopterus lumpus* |

| | | | | | |
|---|---|---|---|---|---|
| 29/05/2024 | Inntian | SES | 52 | 55 | *Cyclopterus lumpus* |

**Table S4. Camera recordings at the kelp farms in April 2024.** OF=Ocean Forest. SES= Seaweed Solutions.

| Date | Site | In time | Out time | Location | Comment | Camera | Shoot speed | Hours running | N of pictures |
|---|---|---|---|---|---|---|---|---|---|
| 06/04/2024 | OF | 08:59 | 10:47 | 2 meters depth, beside production line | Marine snow, 3m vis | 2 | 00:00 | 01:48 | 216 |
| 08/04/2024 | OF | 10:25 | 15:44 | 3 meters depth, beside production line | Marine snow, 3m vis | 2 | 00:00 | 05:19 | 638 |
| 09/04/2024 | OF | 12:00 | 14:49 | 4 meters depth, beside production line | Marine snow, 3m vis | 2 | 00:00 | 02:49 | 338 |
| 16/04/2024 | SES | 11:07 | 20:08 | 3 meters, under production and old production line | | 2 | 00:00 | 09:01 | 3246 |

| | | | | | | | | | |
|---|---|---|---|---|---|---|---|---|---|
| 18/04/2024 | SES | 09:24 | 18:42 | 3 meters up to surface | | 2 | 00:00 | 09:18 | 3348 |
| 16/04/2024 | SES | 11:19 | 20:34 | 3 meters depth, beside production and structural line | | 1 | 00:00 | 09:15 | 3330 |
| 18/04/2024 | SES | 09:20 | 17:10 | 4 meters facing deeper water | | 1 | 00:00 | 07:50 | 2820 |
| **TOTAL** | | | | | | | | **21:20** | **13936** |

**Table S5. Camera records of fish occurences at the kelp farms in April 2024.** OF=Ocean Forest. SES= Seaweed Solutions.

| **Site** | **Fish** | **Abundance** | **Time** | **ID** | **Event** | **Depth** | **Date** | **Comment** | **Location** |
|---|---|---|---|---|---|---|---|---|---|
| OF | Saithe | 1 | 10:41 | G0040275 | 1 | 5 m | 08/04/2024 | Not feeding | 3rd block right side |
| OF | Saithe | 2 | 10:41 | G0040276 | 1 | 5 m | 08/04/2024 | Not feeding | 3rd block right side |

| | | | | | | | | | |
|---|---|---|---|---|---|---|---|---|---|
| SES | Saithe | 4 | 11:30 | G0075028 | 2 | 5 m | 18/04/2024 | Not feeding | 1st block on left, looking into farm |
| SES | Saithe | 11 | 11:30 | G0075029 | 2 | 5 m | 18/04/2024 | Not feeding | 1st block on left, looking into farm |
| SES | Saithe | 32 | 11:34 | G0075052 | 2 | 5 m | 18/04/2024 | Not feeding | 1st block on left, looking into farm |
| SES | Saithe | 13 | 11:34 | G0075053 | 2 | 5 m | 18/04/2024 | Not feeding | 1st block on left, looking into farm |
| SES | Saithe | 2 | 12:21 | G0075334 | 3 | 5 m | 18/04/2024 | Not feeding | 1st block on left, looking into farm |
| SES | Saithe | 8 | 12:21 | G0075335 | 3 | 5 m | 18/04/2024 | Not feeding | 1st block on left, looking into farm |

| | | | | | | | | | |
|---|---|---|---|---|---|---|---|---|---|
| SES | Saithe | 3 | 12:22 | G0075336 | 3 | 5 m | 18/04/2024 | Not feeding | 1st block on left, looking into farm |
| SES | Saithe | 5 | 12:22 | G0075337 | 3 | 5 m | 18/04/2024 | Not feeding | 1st block on left, looking into farm |
| SES | Saithe | 4 | 12:41 | G0075453 | 4 | 5 m | 18/04/2024 | Not feeding | 1st block on left, looking into farm |

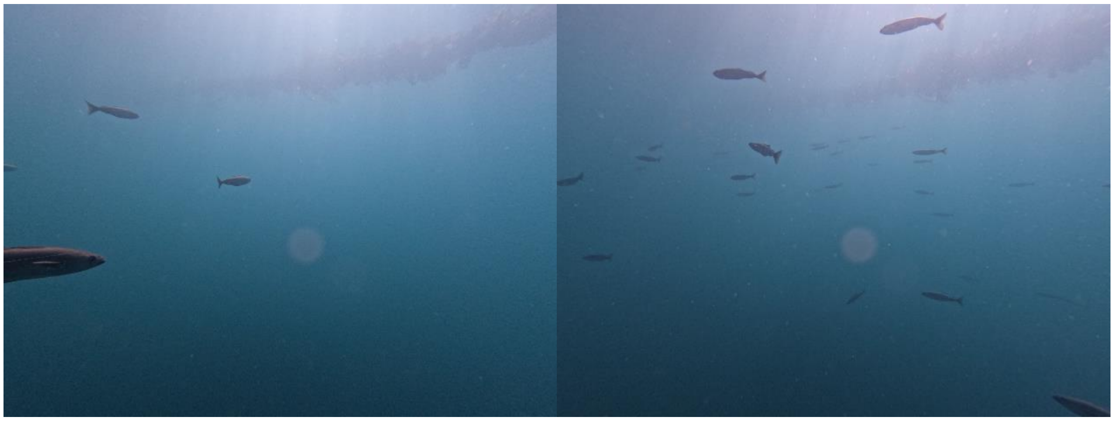

**Figure S1.** Examples of camera observations of fish, showing saithe at the Frøya kelp farm location in schools of up to 32 individuals.

**Fauna observation details from traditional sampling**

Records and details of fish and invertebrates recovered during sampling in 2024 are available in the attached file Supplementary File 1.

**Additional methods to observe fishes at the kelp farm**

Three fish/macro invertebrate traps (94 x 64 x 63 cm) with 25 mm nylon mesh sizes and a 5cm diameter opening were deployed at 2 meter depth at the Austevoll kelp farm site for a duration of 4 days and 4 nights. These were baited with boiled Pandalus borealis, a local shrimp species, in an attempt to catch benthic scavenging fish or invertebrates. The traps resulted in no catches.

One trammel net (30 m long,1.8 m deep, smallest, and inner mesh size 52 mm) was deployed at the Frøya kelp farm site for a duration of 4 days and 4 nights (see details in Table S1). The net was hung from kelp farm structures, placed at a depth of ca. 5 m in an attempt to capture larger fish or fish dwelling at a deeper-depth, if present. The net resulted in no catches.

Camera surveys of 2024 were repeated at the Austevoll kelp farm site in 2025 to confirm that our observations were representative. Thus, two cameras were deployed at both kelp farms to capture fish diversity and behaviour during daylight hours using GoPro Hero 10 Black cameras, recording a photo every 10 seconds. Dates under observation were 01.04.2025 and 27.04.2025. A total of 10 hours and 54 minutes were observed, recording 3,981 pictures that revealed one juvenile lumpfish and 58 images with 1-2 saithe over a single period of 3 hours and 54 minutes (Figure SX).

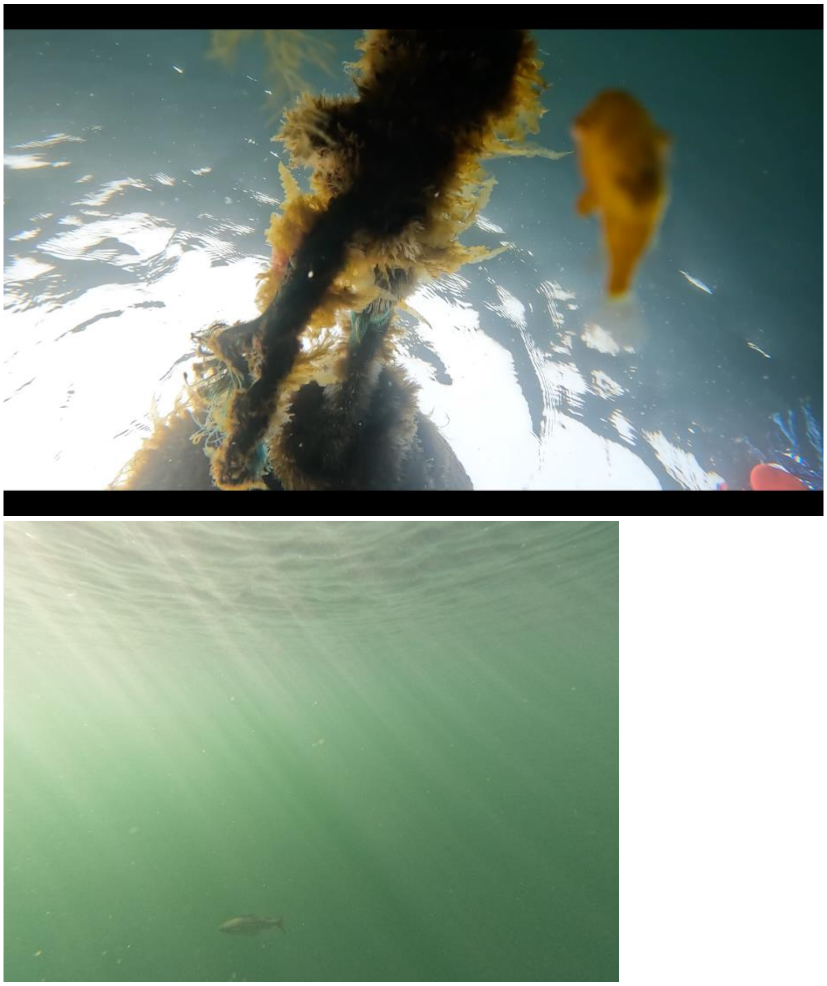

Figure S2. Repeat camera recordings at the  kelp farm site in April 2025, showing a single juvenile lumpfish (left) and saithe (right).

In addition, the hand-capture of fishes at harvest was repeated for the same reason. Approximately 40,000 tonnes was harvested between two sites at Austevoll, one Trollsøy (as in the 2024 survey) and the nearby Flatøysundet kelp farm. Dates of harvest, location and counts of lumpfish (the only species observed) are listed in Appendix Table X, suggesting that 2024 observations are representative. The density of lumpfish cannot be reliably estimated due to sampling being performed ad-hoc.

Table S6. Counts of lumpfish retrieved in 2025 at Austevoll kelp farms during harvesting by the operator Ocean Forest

| **Date** | **Location** | **Count of Cyclopterus lumpus** |
|---|---|---|
| 24.4.2025 | Flatøysundet | 1 |
| 27.4.2025 | Trollsøy | 10 |
| 26.4.2025 | Trollsøy | 14 |
| 26.4.2025 | Flatøysundet | 9 |
| 29.4.2025 | Flatøysundet | 2 |
